# 4D imaging reveals stage dependent random and directed cell motion during somite morphogenesis

**DOI:** 10.1101/280883

**Authors:** James McColl, Gi Fay Mok, Anna H Lippert, Aleks Ponjavic, Leila Muresan, Andrea Münsterberg

**Affiliations:** School of Biological Sciences, Cell and Developmental Biology, University of East Anglia, Norwich Research Park, Norwich, NR4 7TJ, UK; Chemistry Department, University of Cambridge, Lensfield Road, Cambridge, CB2 1EW, UK; Cambridge Advanced Imaging Centre (CAIC), Downing Street, Cambridge, CB2 3DY, UK

**Author notes:** corresponding authors: James McColl, Andrea Münsterberg. Equal contribution/joint first authors (James McColl, Gi Fay Mok).

## Abstract

Somites are paired embryonic segments that form in a regular sequence from unsegmented mesoderm during vertebrate development. Of fundamental importance, they are transient structures that generate cell lineages of the musculoskeletal system in the trunk such as cartilage, tendon, bone, endothelial cells and skeletal muscle. Surprisingly, very little is known about the morphological transition and cellular dynamics during somite differentiation. Here, we address this by examining cellular rearrangements and morphogenesis in differentiating somites using live multi photon imaging of GFP-transgenic chick embryos. We specifically focussed on the dynamic changes in two principle regions within the somite (the medial and lateral domains) to investigate extensive morphological changes. Furthermore, by using quantitative analysis and cell tracking, we were able to capture for the first time a progenitor cell bulk movement towards the rostral-medial domain of the myotome, where skeletal muscle formation first initiates.

## Introduction

Embryonic morphogenesis involves dramatic tissue deformation and growth, which often occurs rapidly over short time-scales. It is implicit that tissue deformations are driven by local cellular activities, including cell proliferation, changes in morphology and/or size, and cell rearrangements. However, it has been challenging to image, capture and quantify these processes in live tissues.

Somites are transient epithelial, near spherical, structures that form during vertebrate development from the presomitic mesoderm (PSM) in a regular sequence and with a rostro-caudal progression (Benazeraf & Pourquie, 2013). As they differentiate, these paired body segments dissociate ventrally where epithelial-to-mesenchymal transition (EMT) leads to formation of the sclerotome, the source of the axial skeleton. The dorsal somite remains epithelial and produces the dermomyotome and myotome, the source of all trunk and limb skeletal muscles (Christ & Scaal, 2008). Signalling and genetic control of this process is well characterised (Buckingham & Rigby, 2014), however surprisingly very little is known about how individual cell dynamics and cellular rearrangements drive morphogenesis within the somite during its differentiation. Therefore, improved and greater understanding of these processes will benefit the derivation of musculoskeletal lineages from pluripotent stem cells (Chal & Pourquie, 2017).

Along the anterior-posterior axis, each individual somite is flanked by neighbouring somites; other adjacent tissues on the medial, lateral and dorsal sides are the neural tube (future spinal cord), the intermediate and lateral plate mesoderm and the surface ectoderm respectively. Signalling molecules derived from these tissues govern the specification of somite cells towards particular fates (Pourquie et al., 1993; Johnson et al., 1994; Fan & Tessier-Lavigne, 1994; Fan et al., 1995; Munsterberg et al., 1995; Munsterberg & Lassar, 1995; Pourquie et al., 1995; Pourquie et al., 1996; Reshef & Lassar., 1998; Kahane et al., 2001; Gros et al., 2004). In addition, these tissues impose rigidity and mechanical constraints, which may contribute to somite morphogenesis. What remains unknown is what morphological processes occur during somite development.

The medial domain of the somite, closest to and running parallel to the neural tube, is particularly important for the formation of skeletal muscle. It is here that the early myotome first forms from early myoblasts, which navigates under the epithelial dermomyotome, and subsequently from cells that enter the myotome from all edges of the dermomyotome. This process has been extensively characterized using intricate cell labelling (Denetclaw & Ordahl, 2000; Kahane et al., 2001; Ben-Yair et al., 2003; Gros et al., 2004; Rios et al., 2011).

Cell proliferation within the dermomyotome contributes to its growth and, in addition, mitotic cells have been observed in the dermomyotome lip (Kahane et al., 2001, Venters & Ordahl, 2005). In epithelial somites, most cells were labelled following a short pulse of BrdU which indicates replication, with exception of some cells abutting the neural tube (Kahane et al., 1998). However, the potential contribution of localized cell proliferation to cell rearrangements, movement and thus to somite morphogenesis has not yet been evaluated.

In addition, the role of force and cell flow is also important for somite development. Discreet patches of laminin within the extracellular matrix (ECM) for example have been shown to form during early somite development. As the matrix develops it provides activating and constraining force regions (Thorsteinsdottir et al., 2011; Martins et al., 2009). Cell movement and cell expansion will generate forces which will push, pull and direct the position of cells and therefore shape tissues. This creates tension and stress responses from cells and surrounding matrices, and with surface tension thought to play a small role in development (Grima & Schnell, 2007) direct force transfer between cells is most likely a primary mechanism.

This work sought to clarify and quantitate the behaviour of individual cells in real time, in a living, developing somite. Thus far our insights regarding the dynamic cell re-arrangements during somite development are limited as almost everything has been derived from the analysis of fixed samples. Using multi-photon time-lapse microscopy and quantitative analyses, we measure somite growth and establish that it is mainly driven by cell proliferation, which is not uniform and instead asymmetrically distributed in a somite. We show that at early and intermediate stages, medial somite regions contain fewer cells and a population of larger cells. For the first time, in living tissue it has been possible to determine that morphogenetic changes are most pronounced in intermediate stage somites. Furthermore, these changes correlate with an increase in proliferating cells. Our in depth analysis also revealed that cells undergo a collective, directed motion towards the emerging dermomyotome lip, where myogenic cells are known to enter the myotome. Taken together, we identify cellular drivers of somite morphogenesis and propose a model whereby the nascent dermomyotome lip creates a sink towards which the cells move due to constraints imposed by surrounding tissues.

## Results

### Somite growth rate

To determine the extent to which migration, proliferation and cell growth contribute to somite growth and changes in tissue shape, we sought to image somite growth directly. By using multi-photon microscopy, we determined the spatiotemporal distribution of cells and their behaviour. Multiple somites were excised intact and oriented together with intact flanking tissues in humidified imaging chambers (Fig. 1 A, B). We imaged the four newest somites formed in the posterior tail and examined their change in volume overall, and separately measured the change in size of the hollow centre (somitocoel) (Fig. 1C, D). Firstly, the growth rate of individual somites was determined over 180 minutes; all somites increased in volume with a rate of 19 +/-4 µm^3^ hr^-1^ (Fig. 1D). In contrast, the size of the somitocoel remained constant (126 +/-4 µm^3^ at time 0 and 124 +/-4 µm^3^ at 180 minutes). Based on this data, we modelled somite growth as a collection of spherical structures to determine the number of cell divisions required if growth is solely dependent on proliferation (Fig. 1D). For a growth increase of 19 µm^3^ hr^-1^ the division rate of somite cells would need to be 3 new cells hr^-1^ for an average cell size of 6 µm (Schindelin et al., 2012). Alternatively, we also propose that the volume change could also be due to increase in cell size, or a combination of both proliferation and size increase (Fig. 1E). Therefore, we tested this hypothesis to identify the number of proliferating cells during somite development and where precisely these cells are located.

**Figure 1:**
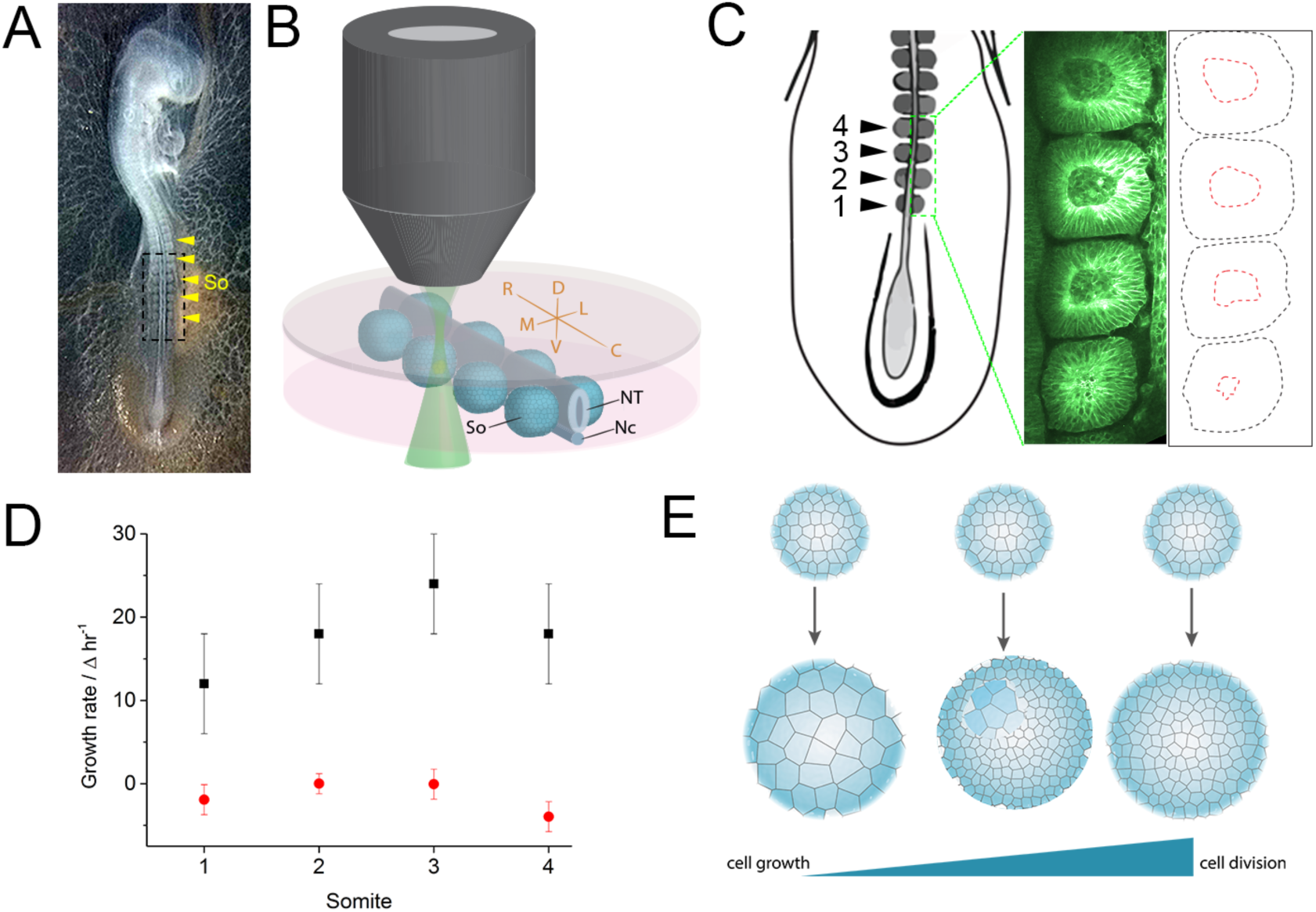
Imaging and measuring somite growth. Somites from 2 day-old memGFP transgenic chicken embryos were dissected as indicated by black dotted line box (A). Somites were positioned and set into an agarose:media gel in chambers and imaged (B). The volume of the four newest formed somites, both the whole somite (black dotted lines) and the somitocoel (red dotted lines), were measured and calculated (C and D, n=4). The growth rates for individual somites were calculated and plotted with an average growth rate of 19 +/-4 µm^3^ hr^-1^ determined for the somite (D). Growth can be modelled as either individual cell growth, cell division or a combination of both (E). So, somites; NT, neural tube; Nc, notochord. Errors are SD.

Gross somite morphological changes along the embryonic axis are development indicators, with more recently formed, posterior somites less differentiated (early stage) compared to more anterior somites (late stage) (Fig. 2A and movie V1-3). We investigated if the rate of change in somite morphology was different for somites as they progress in their differentiation. To do this we compared somites from early, mid and late stages along the embryonic axis; for each stage we determined the change in somite morphology over 180 minutes (Fig. 2B). In addition, we determined the distribution of morphology changes through the tissue by comparing slices from the dorsal, centre and ventral sections of individual somites from early, mid and late stages. By calculating the correlation coefficient for each somite we are able to calculate how much the somite changes with a higher correlation coefficient meaning less change. Interestingly, we found that morphology changes were not linear and mid stage somites changed greatest, in all regions of the somite: dorsal, centre and ventral. By contrast, the early and late stage somites changed at a slower rate with greater change observed in the dorsal region of early somites (Fig. 2B). Thus, mid stage somites were more dissimilar throughout the whole somite, indicating a greater rate of change during the time window measured.

**Figure 2:**
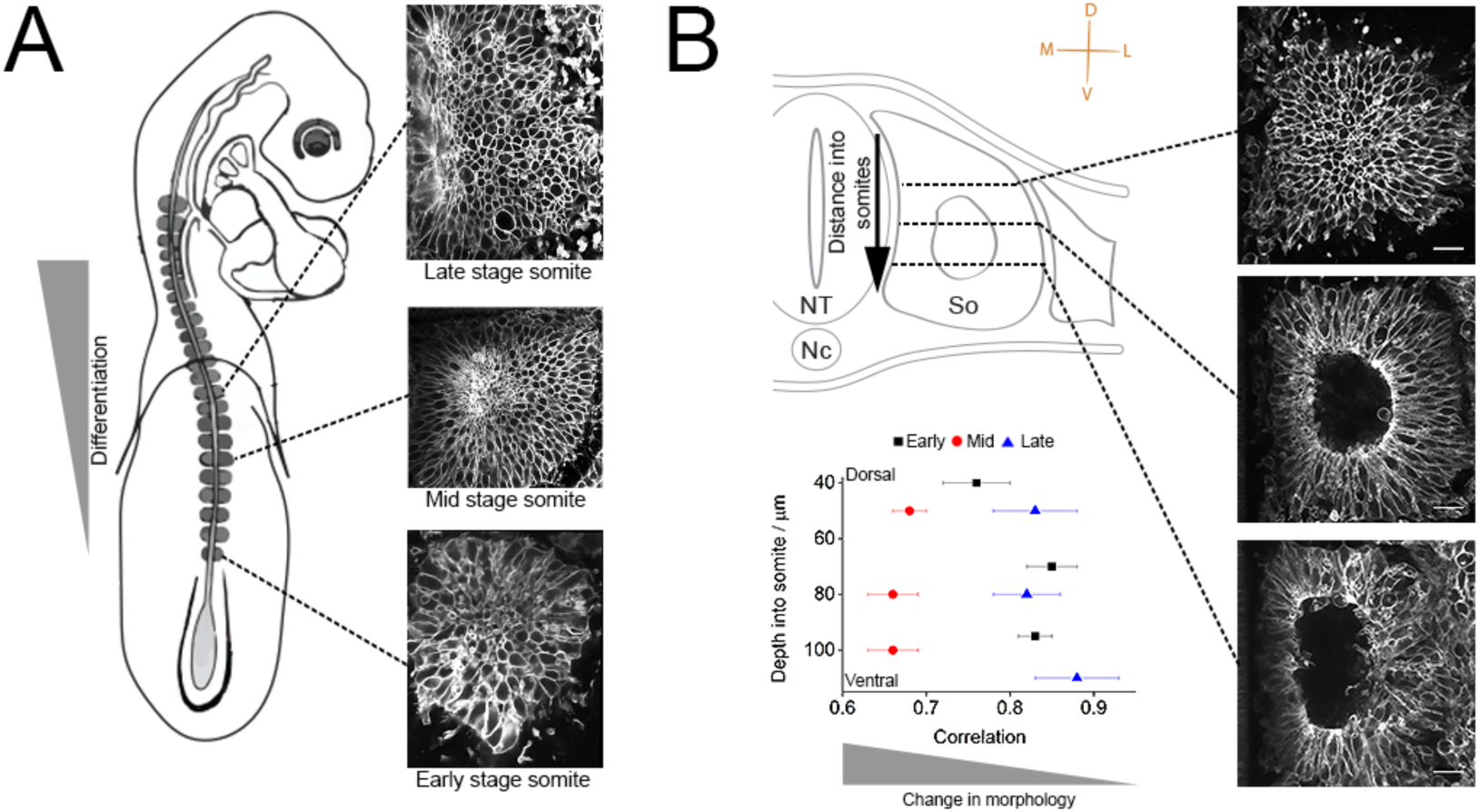
Largest morphological changes are at the mid stage. Early, mid and late stage somites were individually imaged and analysed (A). Top, middle and bottom of early (black square, n=3), mid (red circle, n=4) and late (blue triangle, n=3) stage somites were correlated by overlaying successive images between t0 and t180 min (B). A higher correlation value indicates less change over time. Scale bars 10µm.

### Morphological changes at the medial and central domain

Signals from adjacent tissues, including the surface ectoderm, neural tube and migrating neural crest cells are pivotal for cell specification within somites (Fan & Tessier-Lavigne, 1994; Munsterberg et al., 1995; Borycki et al., 1999; Rios et al., 2011). This includes Wnt and Notch signals, which instruct medial somite progenitor cells to move from the medial region into the somite (Siero et al., 2016). To determine the behaviour of cells in the medial against the lateral domains within somites we measured a range of parameters including the number and size of cells over time in the medial domain (MD) (the nascent dermomyotome lip) and compared this to the lateral domain (LD) (Fig. 3A). The number of cells at the medial domain is similar for both early and mid-stages, 81 +/-12 and 68 +/-3, respectively. At the LD, a higher number of cells was found in mid-stage somites, 212 +/-8, when compared to early stage somites, 160 +/-18 (Fig. 3A). When examining cell size, we found a more uniformly sized cell population at the LD compared to the MD where cell size varied greatly in both early and mid-stage somites (Fig. 3A). Quantification of cell size distribution revealed that there are two populations with a discrete size difference in the medial domain (Fig. 3B). In addition, we assessed the number of splitting events by calculating the large rounded cells. We find that the splitting rate was much greater in mid stage somites than in early or late stage somites, (p value = 4.7e^-8^, for mid against early and p value = 1.3e^-4^, for mid against late, two tailed t-test). The early and late populations were not significantly different (p value = 0.2) (Fig. 3C). This is consistent with the observation that mid stage somites had a greater rate of change.

**Figure 3:**
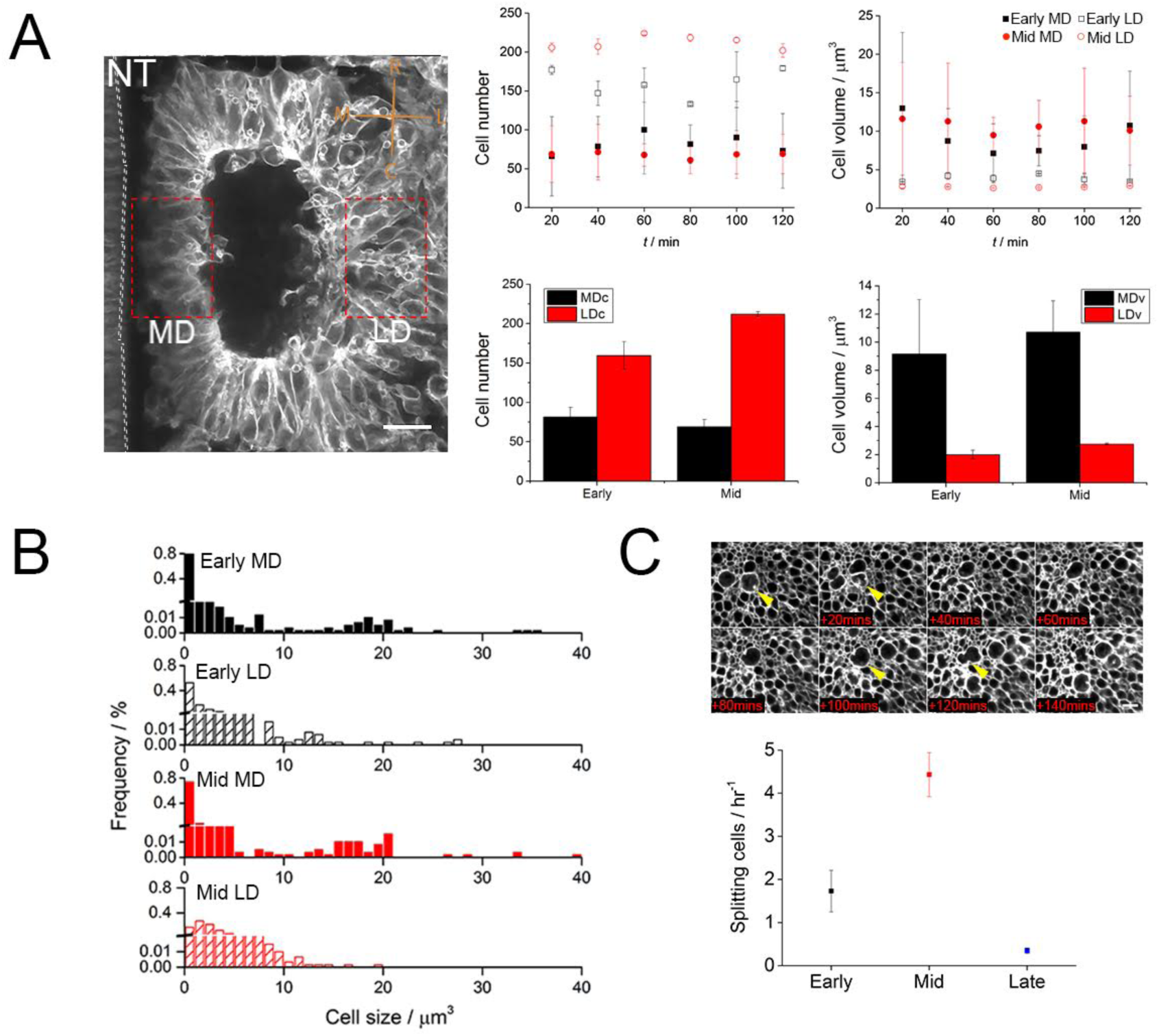
Cell size and number are heterogeneous across the somite. Cell number and volume were determined over time for early (black square) and mid (red circle) stage somites at the medial domain (MD) and lateral domain (LD). Average comparisons for all data sets, MDc and LDc denote medial and lateral cell number. MDv and LDv denote medial and lateral cell number (A). Cell size histograms for MD (solid fill) and LD (pattern fill) for early and mid-stage somites (B). Cell proliferation rate was determined by counting splitting cells (yellow arrow head). Cell proliferation at the LD and proliferation rates for early (black, n=3), mid (red, n=4) and late (blue, n=3) stages were measured. Errors are SD and Scale bar 20µm.

### Concerted movement of cells towards rostral medial regions

To further understand the mechanisms underlying the greater rate of change in mid-stage somites, we characterized cell behaviours in more detail. In the rostral-medial corner we find detaching cells, which suggests an epithelial to mesenchymal transition in this region (Fig. 4A). To determine the cellular processes that may contribute we tracked individual cell movement in the dermomyotome region, in mid as well as early and late stage somites (i.e. above the somitocoel or above the ventral mesenchymal part of the somite respectively) (Fig 4B and movies V4-6). Using ImageJ and custom written Matlab software, we determined both the direction (Fig. 4C, D) and speed (Fig. 4E) of cells. In early-stage somites, movement of cells appeared random, with a velocity of 0.14 ± 0.08 µm min^-1^ (Fig. 4C-E). Interestingly, for mid-stage somites, we observed a concerted movement of cells directed towards the rostral-medial quadrant (Fig. 4C-D). By plotting the angle of the tracks as a function of their length, we find that the longest tracks are both directed and centred around the mean tracks angle for mid-stage somites (Fig. 4D). We also note that more cells were moving faster in mid-stage somites 0.21 ± 0.11 µm min^-1^ (Fig. 4E). Plotting the mean direction of the tracks gives the overall directionality (Fig. 4F), which was higher for mid-stage somites compared to early and late stages. In late-stage somites we observed cells moving inwards, towards the centre of the maturing somite, at a slower speed of 0.06 ± 0.03 µm min^-1^ (Fig. 4C-E). Overall our measurements revealed a directed and concerted cellular motion towards the rostral-medial region of the mid stage somite, as well as suggesting a slower motion towards the centre of the dermomyotome at a later stage (Fig. 4G).

**Figure 4:**
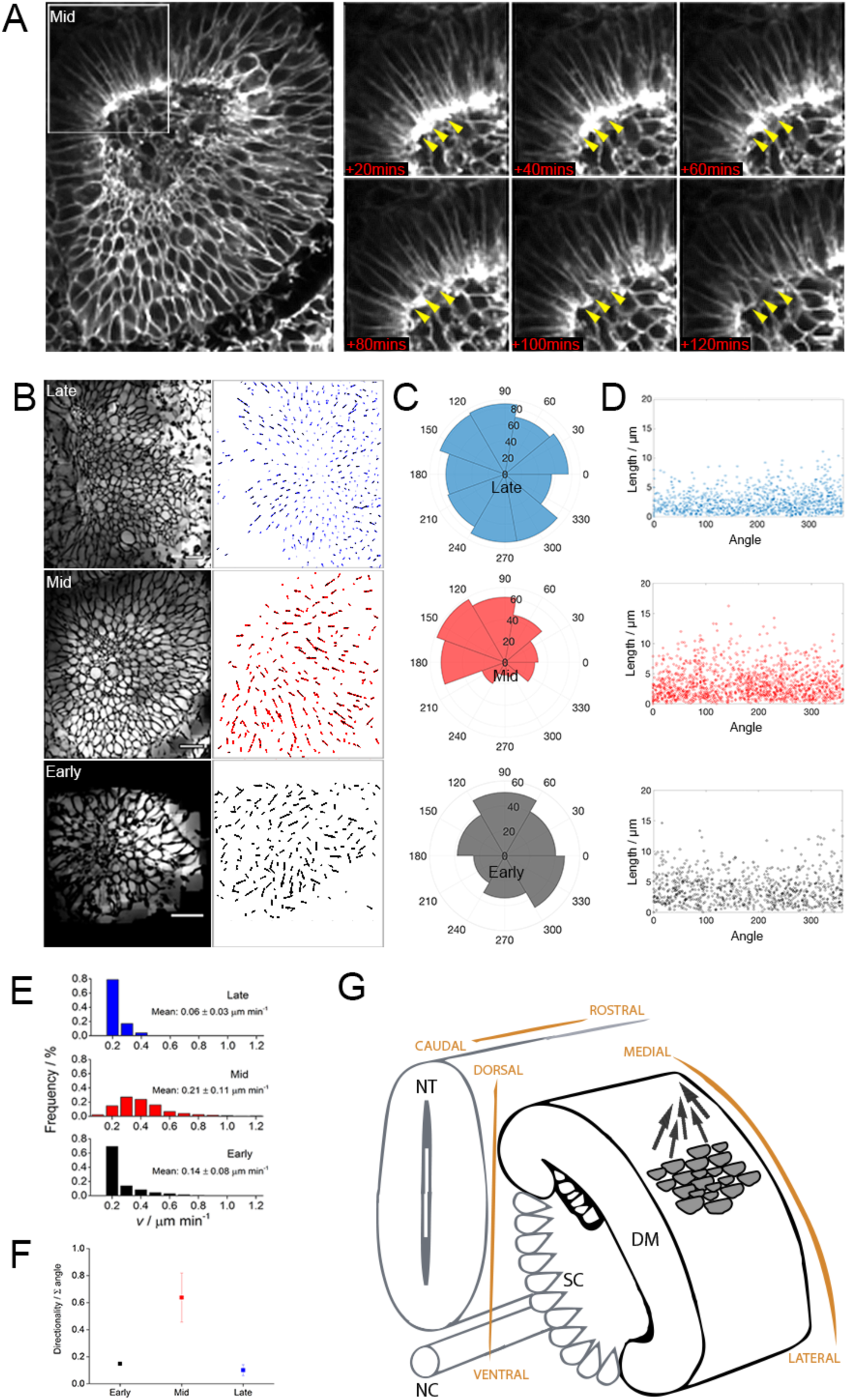
Directional concerted cell movement in mid-stage somites towards the rostro-medial region where individual cells dissociate (arrows). Scale bar is 10µm (A). Inverted images are used to track individual cells. Scale bars 20µm (B). Angle distribution for individual tracks of early (black, n=3), mid (red, n=4) and late (blue, n=3) somites (C). Length versus angle plots for all tracks (D). Track speed for early (black), mid (red) and late (blue) somites measured (E). Summing all tracks gives directionality for early (black), mid (red) and late (blue) stages (F). Directional concerted cell movement model for mid stage somites (G). DM, dermomyotome; NT, neural tube; NC, notochord; SC, sclerotome.

## Discussion

Somite differentiation is a dynamic process, during the elongation of the vertebrate embryonic axis, involving rapid development in size and shape. Here we used real-time multi-photon imaging combined with precise quantitative analysis of cell size and number, in addition to detailed observations of localized cell behaviours to characterize this important process. This has provided surprising insights into cellular mechanisms underlying somite morphogenesis.

We used an ex vivo system with multiple somites surrounded by other native tissues, which are known to provide extrinsic signals and inductive cues (Fan & Tessier-Lavigne, 1994; Munsterberg et al., 1995; Borycki et al., 1999; Rios et al., 2011). The tissues maintained their structural integrity with viable cells that displayed coordinated behaviours and extended filopodia (not shown). We observed cell divisions and there was no indication of apoptosis. Furthermore, we observed in real time the morphological changes anticipated based on what is known from fixed whole embryo samples (Kaehn et al., 1988; Kahane et al., 1998; Kahane et al., 2002). For example, late stage somites formed a myotome layer (Fig. 2A and movie V1-3) as has been described previously (Gros et al., 2004; Rios et al., 2011). Thus, we believe that our quantitative ex vivo measurements closely reflect what is occurring in intact embryos.

Somites grow in size as they develop over a short period of time (Fig. 1D) and this growth could be due to an increase in cell size, cell proliferation or a combination of both (Fig. 1E). We determined somite growth rate of 19 µm^3^ min^-1^ for early somites, which equates to an increase of approximately 3 cells hr^-1^ if the growth is due to proliferation alone. We found that the size and number of cells increased with time, however interestingly greatest size increase occurred predominantly at the medial domain (MD) (Fig. 4C, D). The contribution of proliferation for early somites is 1.7 cells hr^-1^ (Fig. 3C), suggesting that approximately half of somite growth is achieved through proliferation and half through growth of individual cells at early stages (Fig. 1E). Furthermore, we observe larger cells at both early and mid-stage MDs suggesting that the majority of growth at the mid stage comes from proliferation. Proliferation rates increased for mid stage somites to 4.4 cells hr^-1^ and we observed that this mainly occurred at the lateral domain (LD) of the somite. Greater cell numbers and increased proliferation in the LD could be responsible for the directed motion we observe of cells from lateral towards rostro-medial (Fig. 4A, B, G and movies V4-6). For late stage somites we found a low proliferation rate of 0.3 cells hr^-1^ and observed that these cells moved slowly, suggesting somite growth had almost stopped. It is likely that other processes such as differentiation of the myotome and muscle fibres begin to dominate at this stage (Fig. 3C, 4E).

We next sought to determine the distribution of proliferating cells across the somite to underpin the notion of somite growth is contributed by proliferation and locate the regions proliferation was greatest. Using real-time observation, we find that the cell number at the MD remained relatively constant at early and mid-stages, whereas the number of cells at the LD increased significantly (Fig. 3A). This non-homogenous distribution is in agreement with post-mitotic cells observed using BrdU pulse labelling in the medial aspect of epithelial somites (Kahane et al., 1998).

To study further in mid-stage somites, we observed whole somite morphological changes were more pronounced in mid-stage somites and changes in cell number and size significantly contributed to this. Although early stage somites underwent less morphological change it is interesting that the dorsal region of the somite changed more and had a lower correlation coefficient (Fig. 2B). Based on the direction of tracked individual cells their motion appears random and we further observe that the longest tracks for this stage radiate across the somite in various directions (Fig. 4C-D). This behaviour could be explained if individual or small populations of cells move into different regions, as has been previously proposed (Kahane et al., 1998), however, we could not observe this directly. By contrast, for mid-stage somites we identified a directed movement of dermomyotomal cells towards the rostral-medial domain, where cells were detaching (Fig. 4A). The longest tracks were concentrated in a single direction (Fig. 4B-D). Desmin-positive myogenic cells are present in the rostro-medial, mid-stage somite (Kaehn et al., 1988; Kahane et al., 2002), and we propose that the collective motion of dermomyotome cells towards this domain may contribute to the fate decisions of these progenitors, for example by providing a mechanical stimulus as has been observed in stem cell differentiation (McBeath et al., 2004; Delaine-Smith, 2012; Pattie, et al., 2012).

The directed cell motion we observe originates predominately from the caudal regions and we propose that it is moving cells towards a sink at the rostral– medial domain. Although outside the scope of this work, it is likely that multiple factors are responsible for how these cells are able to orchestrate a precise motion towards a sink. Cells in the LD region may receive signals, including Wnt signals, from the overlying ectoderm (Sagar et al., 2015). Medial lip cells have been shown to migrate via the transition zone into the centre of the somite after receiving signals from the neural crest (Rios et al., 2011). Ingressing cells at the MD are also some distance from the proliferating cells (predominantly LD), suggesting that they may not be involved in the concerted cell movement identified here, although they may still play a role in EMT. This novel observation of a concerted motion of cells towards the rostral-medial domain of mid-stage somites, where cells are detaching, is exciting as it correlates with the presence of early myoblasts at the rostro-medial corner of the dissociating somite (Kaehn et al., 1988; Kahane et al., 2002). We speculate that this cellular motion, concomitant with EMT, may in fact contribute to the appearance of early myoblasts. In addition, we propose that asymmetrically distributed cell proliferation events, predominantly in the lateral region of the somite, are likely to play a key role by creating local forces and tension. We suggest that some of the cellular mechanisms revealed here, may be of more general importance for the shaping of tissues during embryonic development.

## Materials and Methods

### Embryo culture, dissection and mounting

Fertile eggs, transgenic for membrane-bound GFP (MemGFP, Roslin Institute, University of Edinburgh, UK) were incubated at 37°C in a humidified incubator until HH14/15. Embryos were harvested in Glutamax Ham’s F12 media (cat. no. 31765) and explants containing somites, intermediate mesoderm, surface ectoderm, endoderm and neural tube were dissected using fine scalpels. We prepared low melting point agarose (Invitrogen cat. no. 16520-050, 2% w/v) in Glutamax Ham F12 media, then added 10% Fetal bovine serum and 1% penicillin/streptomycin after the agarose/media had cooled. The dissected tissue was positioned with the dorsal side towards the base of the glass slide, on non-coated 35 mm MatTek dishes with 7mm glass diameter (P35G-0-7-C) and warm agarose/media added to the somite explants until fully covered. Dishes were placed on ice for 5 minutes before placing inside a humidified heat chamber (37°C) for microscopy.

### Long-term two-photon microscopy

Individual somites were imaged as stacks of slices with each slice being 1000 x 1000 pixels (X=250nm Y=250nm Z=510nm). This enabled us to image an individual somite in 20 minute intervals, which included a 5 minute rest period. The number of slices was determined by the size of the somite at 20x magnification using a TriM Scope II (Labtec instruments) two-photon microscope. Initial laser power was increased exponentially through the stack to allow for signal loss due to deeper imaging. Low magnification images were recorded at 10x magnification.

### Tracking and image analysis

Images were first denoised using **nd-safir,** an image sequence denoising software to improve the signal to noise ratio of the images (Boulanger et al.,2010). Following this, somites were aligned in Z manually and slices through the somite over time were selected using custom written Matlab code.

To determine the correlation coefficient images at time 0 and time 180 minutes were compared and correlated using Image_correlator plugin from ImageJ. Here, two images are compared pixel by pixel to determine their correlation. The value for each pixel is plotted as X for one image and Y for the other and once every pixel value is plotted the fit for the plots determines the correlation. If the two images are identical, the fit would be 1.

To track the cells, time slice stacks were inverted, bandpass filtered and background subtracted using a rolling ball radius of 20 pixels (ImageJ). Tracking was performed using the ImageJ trackmate plugin, simple LAP tracker (Schindelin et al., 2012). We then use custom written matlab code to calculate the average direction for each track and it is these values which we use to plot the radial distribution and length versus angle plots. Summing all directionality values indicate how directed the cells were for early, mid and late stages. If the directions are random the directionality would be low as tracks moving in opposite directions would cancel out.

### Low mag whole somite size quantification

Somites were modelled as spheres over time by taking the central image of the somite and determining the perimeter using ImageJ. From the perimeter, we calculate the area of the somite as a sphere.

### 3D MD and LD quantification

Identical XY and Z regions from the medial domain (MD) and lateral domain (LD) were analysed on denoised somites. The Real-time Accurate Cell-shape Extractor (RACE) analysis software was used to determine all cell parameters (Stegmaier et al., 2016) with custom written Matlab code used to extract and plot the data. To determine the cell parameters within the two fixed regions at the MD and LD of early and mid-stage somites we used the RACE algorithm and compared the two regions over time.

### Quantification of cell proliferation

The number of rounded cells were counted using the object detector in ImageJ. Rounded cells were detected by searching for large objects with high circularity in one dataset to determine the ideal paramaters semi-quantitatively. Parameters of minimum diameter 12 µm and 0.85 circularity were then applied to all data setzs. The process of cell division (splitting) takes around 25min, given that we used intervals of 20 minutes this should ensure accurate tracking.

### Statistical methods

To determine significance, we use two-tail distribution T tests with unequal variance.

## Acknowledgements

We thank Dr Paul Thomas, Director of the Henry Wellcome Laboratory for Life Cell Imaging, for advice and support, and Dr Tim Grocott for discussions. JMC was supported by BHF Project Grant (PG/11/118/29292) and GFM by BBSRC grant (BB/N007034/1) to AM.

